# Social and Contextual Memory Impairments Induced by Amyloid-β Oligomers are Rescued by Sigma-1 Receptor Activation

**DOI:** 10.1101/2024.11.16.623839

**Authors:** Souhail Djebari, Raquel Jiménez-Herrera, Guillermo Iborra-Lázaro, Lydia Jiménez-Díaz, Juan D. Navarro-López

**Affiliations:** Neurophysiology & Behavior Lab, Institute of Biomedicine and Instituto de Investigación Sanitaria de Castilla-La Mancha, University of Castilla-La Mancha, Ciudad Real, Spain

**Keywords:** Alzheimer’s disease, amyloid-β oligomers, Social memory, Spatial memory, hippocampus, sigma-1, PRE-084, LTP

## Abstract

Sigma-1 receptors (S1Rs) are widely expressed throughout the central nervous system and modulate neuron intracellular calcium levels, leading to changes in neurotransmitter release and neuronal activity. They also interact with various proteins and signaling pathways, playing a key role in regulating synaptic plasticity in brain areas such as the hippocampus, thereby influencing learning and memory processes. This opens a research avenue to explore S1R modulation as a potential therapeutic target in diseases involving hippocampal synaptic alterations and compromised cognitive processes, such as Alzheimer’s disease (AD). Here, we hypothesize that pharmacological activation of S1R could counteract synaptic plasticity deficits and hippocampal-dependent cognitive alterations in an early-stage amyloidosis model of Alzheimer’s disease, induced by intracerebroventricular (*icv*) administration of A*β*_1-42_ oligomers (oA*β*_1-42_). For that purpose, we investigate *ex vivo* CA3-CA1 synaptic plasticity, while *in vivo*, we performed open field habituation and social recognition tasks to assess contextual and social memory, respectively. Our data show that pharmacological activation of S1Rs with the selective agonist PRE-084 counteract oA*β*_1-42_ deleterious effects on CA3-CA1 long-term synaptic plasticity (LTP), and hippocampal-dependent contextual and social memory, without alterations of spontaneous behaviors. Together, these results provide evidence for the role of S1Rs in ameliorating hippocampal synaptic and contextual memory dysfunctions and, for the first time, in early amyloid-induced social memory deficits, highlighting their potential in the development of comprehensive treatments for early AD. Also, the absence of adverse behavioral outcomes associated with PRE-084 treatment accentuates its safety profile, underscoring its potential as a therapeutic agent.

## 1. Introduction

Alzheimer’s disease (AD) is characterized by a progressive cognitive decline, with its initial pathogenesis strongly associated with the detrimental impacts of the amyloid-*β* (*Aβ*) peptide, particularly *Aβ*_1-42_ [1]. In its early stages, soluble oligomeric forms of *Aβ (*o*Aβ*) increases neuronal activity [2], predominantly within the hippocampus, disrupting the critical balance necessary for synaptic integrity and cognitive function [3, 4]. Deeping our understanding of these initial aberrations is pivotal for clarifying the early cognitive deterioration in AD and identifying potential therapeutic targets to slow its progression. This pursuit has motivated the development of animal models that replicate early neuronal and network anomalies [5] or focus on early hippocampal amyloidosis, revealing the typical detrimental effects of *Aβ*_1-42_ oligomers (o*Aβ*_1-42_) [6]. Within this model, the G-protein-gated inwardly rectifying potassium (GIRK) channel has emerged as a potential therapeutic avenue to normalize neural excitability and counteract these deleterious effects [7–9]. However, the intricate nature of amyloid pathology requires the exploration of additional, complementary targets to enhance therapeutic efficacy and safety. In this context, the sigma-1 receptor (S1R), known for its distinctive properties [10], stands out as a promising candidate for further exploration.

S1Rs, once classified as a subtype of opioid receptors, are now acknowledged as a distinct class of intracellular chaperone proteins. They have garnered considerable attention due to their multifaceted functions within the central nervous system [11]. Predominantly situated in the endoplasmic reticulum, S1Rs are instrumental in modulating calcium signaling [12], thereby exerting a broad influence on neurotransmission systems [13, 14]. Their ability to interact with a diverse array of proteins and signaling pathways significantly broadens their functional repertoire. Notably, S1Rs modulate the dynamics of multiple voltage-activated and ligand-activated ion channels, contributing to the regulation of neuronal excitability and the transmission of information within brain circuits [15]. In the hippocampus, their association with key synaptic receptors, including AMPA and NMDA receptors, underscores their crucial role in synaptic plasticity [16–18]. This connection has significant implications for memory and learning processes, as evidenced by the anti-amnesic effects of S1R agonists [10], and allows to propose S1Rs as promising targets for cognitive enhancement and neuroprotection in disorders marked by cognitive deficits, including AD.

The critical function of S1Rs in modulating neuronal excitability and synaptic plasticity identifies their modulation as a potentially effective intervention in amyloid-related conditions. By rectifying disrupted excitability levels and restoring hippocampal synaptic functions, S1Rs modulation may alleviate memory and learning deficits characteristic of AD. This study examines deeper, extending the scope of previous research to encompass intricate and less explored aspects of cognitive dysfunction induced by amyloidosis, such as social memory. Importantly, this research incorporates a gender perspective by including mice of both sexes, acknowledging the pivotal role of sex-specific differences in AD pathology and treatment outcomes [6, 19]. This comprehensive exploration enriches our understanding of the therapeutic potential in addressing synaptic dysfunctions during the early stages of AD. Preventing these initial dysfunctions from escalating into extensive neuronal damage is imperative, and S1R may play a key role in this preventive strategy.

## 2. Materials and methods

### 2.1 Animals

21 male and 24 female C57BL/6 mice, aged between 5 and 7 months and with body weights ranging from 25-40 g (bred in-house) were utilized. These mice were housed under controlled environmental conditions: a 12-h light/dark cycle, temperature maintained at 21 ± 1 °C, and humidity at 50 ± 7%. Access to food and water was provided *ad libitum*. Prior to surgery, mice were group-housed with a maximum of 5 animals per cage. Following surgical procedures, individual housing was implemented to facilitate optimal recovery and health monitoring. To mitigate the influence of circadian rhythms, all experimental interventions were conducted at consistent intervals during the day. Additionally, regular handling was employed to reduce stress related to the experimental processes.

Ethical clearance for all experimental protocols was granted by the Ethical Committee for Use of Laboratory Animals at the University of Castilla-La Mancha (approval numbers PR 2021-12-21 and PR-2023-21). These protocols adhered to the guidelines of the European Union (Directive 2010/63/EU) and were in compliance with Spanish regulations for the use of laboratory animals in chronic experiments (RD 53/2013, as published in BOE on 08/02/2013 and updated on 24/02/2021).

### 2.2 Surgical procedure for establishing and early amyloidosis model

The generation of an early amyloidosis model in mice required a surgical operation for chronic drug administration, following established methodologies [6]. Initially, mice were anesthetized using a calibrated R580S vaporizer (RWD Life Science, US) with a 0.5 L/min O_2_ flow. Anesthesia induction involved 4% isoflurane (#13400264, ISOFLO, Proyma S.L., Spain), subsequently maintained at 1.5% isoflurane throughout the surgical procedure. Post-operative analgesia was provided through intramuscular administration of buprenorphine (0.01mg/kg; #062009, BUPRENODALE, Albet, Spain), and Blastoestimulina cream (Almirall, Spain) was applied for wound healing.

For intracerebroventricular (*icv.*) administration of o*Aβ*_1-42_, a 26-G stainless steel guide cannula (Plastics One, US) was precisely implanted into the right ventricle of each mouse (1 mm lateral and 0.5 mm posterior to bregma; depth 1.8 mm from brain surface)[20]. Nissl staining was employed post-implantation to verify the accurate positioning of the cannula (Fig. 1A).

**Fig. 1.**
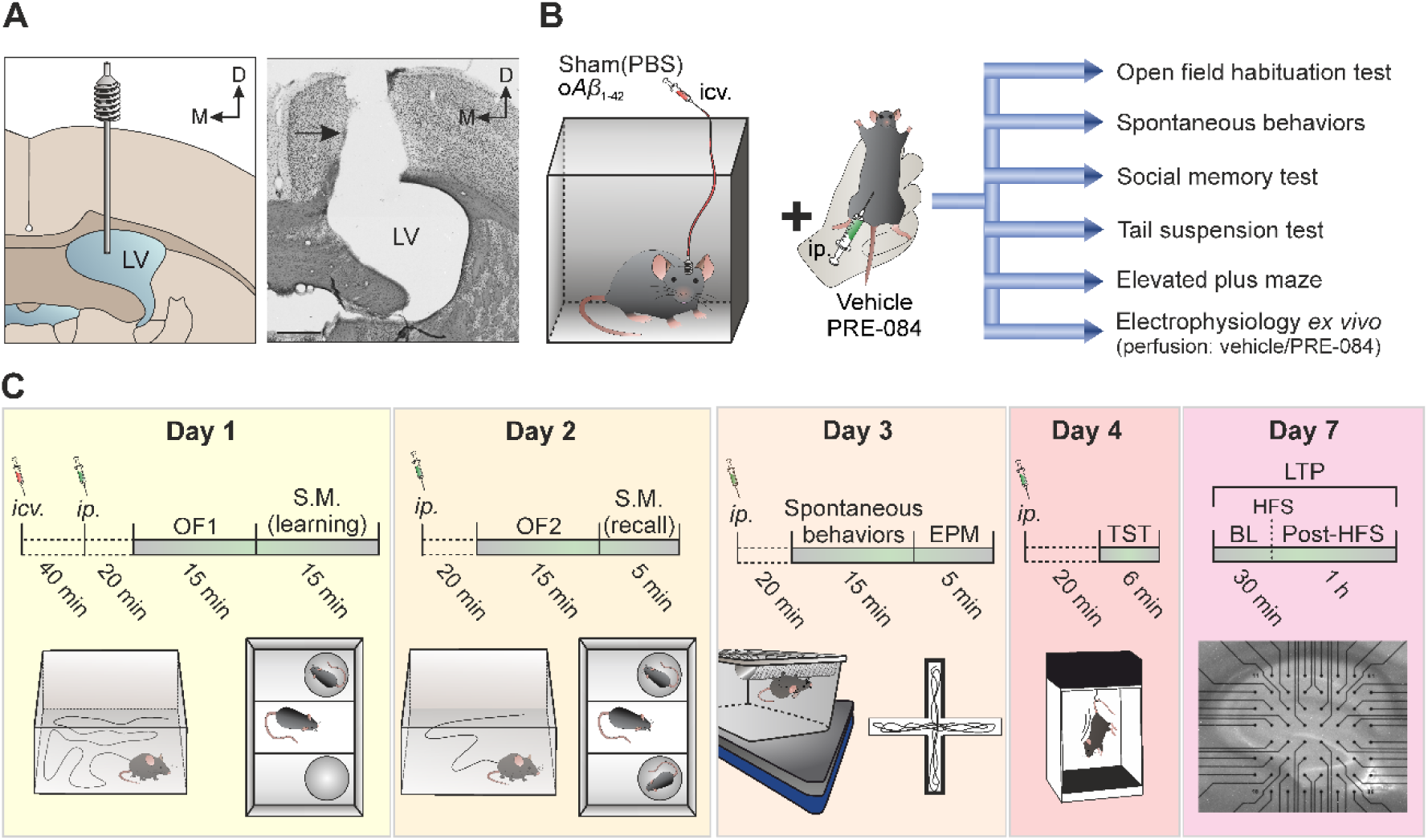
Experimental design and temporal sequence. **A.** For the *icv.* administration of veh. (PBS; sham animals) or o*Aβ*_1-42_, a stainless-steel guide cannula was implanted into the lateral ventricle (LV). The accurate placement of the cannula was histologically verified, as indicated by the black arrow in the right photomicrograph. Scale bar, 500 μm. **B.** Additionally to *icv.* injections, mice received *ip.* injections of veh. (saline) or PRE-084 prior to the behavioral tests. **C.** Temporal sequence of the experiments. Day 1 involved *icv.* injection of veh. or o*Aβ*_1-42_ in freely moving mice and 40 min later *ip.* injections of veh. or PRE-084. Subsequently, the training session of the open field (OF) habituation test and the learning session of the social memory test (S.M) were conducted. On subsequent days (days 2, 3 and 4), one *ip.* injection was administered daily, and, 20 min post-injection, various behavioral tests (or phases of the same test) were conducted sequentially (day 2, recall sessions of the OF and social memory test; Day 3, spontaneous behaviors and elevated plus maze (EPM); Day 4, tail suspension test (TST)). Electrophysiological studies focusing on synaptic plasticity were performed on day 7, utilizing an LTP-inducing protocol of high frequency stimulation (HFS) at the CA3-CA1 dorsal hippocampal synapse. fEPSPs were recorded both before (baseline; BL) and after HFS (post-HFS). *icv.*, intracerebroventricular; *ip.*, intraperitoneal.

A minimum recovery period of one week was observed post-surgery before starting any experimental procedures. Subsequently, mice were randomly assigned into two groups: control sham healthy animals receiving 3 μL of phosphate-buffered saline (PBS) as vehicle (veh.; referred to as sham mice) and mice receiving o*Aβ*_1-42_ (referred to as o*Aβ*_1-42_ mice; Fig. 1B). o*Aβ*_1-42_ (Abcam #ab120301, Cambridge, UK) prepared in PBS as described elsewhere [6, 21, 22], were administered to conscious mice. Administration involved a 3 µL *icv.* injection of o*Aβ*_1-42_ at a concentration of 1 μg/µL, delivered through a Hamilton syringe via an insertion cannula attached to the pre-implanted guide cannula at a rate of 0.5 μL/min [6]. The o*Aβ*_1-42_ *icv.* injection effectively reaches both dorsal hippocampi inducing acute dorso-hippocampal amyloidosis, as verified previously by western blot analysis [6]. This model, representing an early-stage AD-like pathology, has been extensively characterized and validated by our group and others [6–8, 23]

Additionally, 40 min post-*icv.* injections on the initial day and on subsequent days (days 2, 3, and 4), the animals were administered intraperitoneal injections (*ip.*) of either veh. (saline sham+veh._(ip.)_ and o*Aβ*_1-42_+veh._(ip.)_ groups) or PRE-084 (1mg/kg; sham+PRE-084_(ip.)_ and o*Aβ*_1-42_+PRE-084_(ip.)_ groups; Merck #P2607, Darmstadt, Germany) dissolved in saline at a stock concentration of 5.6 mM, 20 min prior to the execution of various behavioral tests (Fig. 1B,C). The timing and frequency of the *ip.* injections were based on established literature [24] while the schedule for the single *icv.* injection was determined based on our previous experiences [6–8].

### 2.3 Open field habituation task

The Open Field (OF) habituation task, a method to assess contextual non-associative learning processes related to hippocampal function, particularly the habituation of exploratory behavior in new environments [25], was employed in this study. This task was initiated 20 min after the *ip.* administration of veh. or PRE-084 and 1 h subsequent to the *icv.* infusion of veh. or o*Aβ*_1-42_ (Fig. 1C). The first training session (OF1), which lasted for 15 min, involved placing the mice at the center of a square acrylic box (base dimensions: 38 x 22.5 x 4 cm; top dimensions: 43.5 x 27.5 x 22.5 cm). Here, they were allowed to explore freely for the session’s duration. On the following day, a second *ip.* injection of veh. or PRE-084 was administered 20 min prior to reintroducing the mice to the same environment for a comparable time span, in a subsequent habituation session (OF2; recall trial).

To monitor and quantify the distances covered by each mouse per min during the OF1 session, as well as to observe alterations in exploratory behavior across sessions, the LABORAS^®^ (Laboratory Animal Behavior Observation Registration and Analysis System; Metris, Netherlands) was utilized [6]. This advanced system translates mechanical vibrations, generated by the animals’ movements, into electrical signals through a sensor platform positioned beneath the cage.

### 2.4 Social memory test

At the conclusion of each open field habituation task session, a social memory test was conducted using a three-chamber cage (40 × 23 × 4 cm for the Plexiglas base arena and 40 × 23 × 11 cm for the top; Fig. 1C). This test also employed the LABORAS®, which records mechanical vibrations resulting from the animals’ movements. Initially, the subject mouse underwent a habituation phase for 5 min in the presence of two empty cylinders placed on opposite sides of the chamber. Following this, the learning session was initiated: the subject mouse was placed in the central chamber, and a novel stimulus mouse of the same sex was introduced into one of the cylinders. During this 10-min session, the subject mouse was allowed to explore freely. The recall session took place 24 h later, wherein the subject mouse was re-introduced to the previously encountered mouse, now a familiar stimulus, and a new novel stimulus mouse of the same sex in a different cylinder for a duration of 5 min. Interaction time was recorded when the subject mouse came within 1 cm of one of the cylinders. To ensure olfactory neutrality, the arena was cleansed with 70% ethanol between each test and allowed to dry completely, effectively removing residual odors.

### 2.5 Elevated plus maze

To assess anxiety-like behavior in mice, the elevated plus maze (LE 842, Panlab, Spain) was utilized on day 3 (Fig. 1C) [26]. This apparatus consists of a cross-shaped platform constructed from methacrylate, comprising two open arms (each measuring 65 x 6 cm) without walls, and two closed arms of identical dimensions but equipped with 15-cm-high opaque walls. The arms are positioned perpendicularly to one another, converging at a central square platform (6.3 x 6.3 cm). The entire structure is raised to a height of 40 cm above ground level.

During the test, each mouse was placed on one of the open arms and allowed a single exploration session of 5 min. The primary indicators for assessing anxiety-like behavior included the number of entries into the open arms and the duration spent within these arms. In addition to these measures, locomotor activity was evaluated based on the number of entries into the closed arms and the total entries, which encompass both open and closed arm entries.

### 2.6 Spontaneous behaviors

On days 3 and 4 post-*icv.* injection, mice underwent a series of behavioral assessments to evaluate their general condition and spontaneous behaviors (Fig. 1C), as previously described [6, 27]. These evaluations included the use of the LABORAS® to analyze stereotypic and locomotive behaviors.

For these assessments, each mouse was placed in a rectangular LABORAS® cage for a 15-min session, employing a methodology previously described. The primary focus was on identifying stereotypic behaviors, with a particular emphasis on grooming, which is often considered an indicator of stress-related behavior. In addition, locomotor activities were quantified, encompassing the frequencies of climbing and rearing behaviors. The LABORAS®, integrated with specialized software (Metris, Netherlands), digitally captured and processed the data from these behavioral tests.

### 2.7 Tail suspension test

To assess depression-like behaviors in mice [28], the tail suspension test was conducted on day 4 (Fig. 1C) using a specialized apparatus (model BIO-TST5, Bioseb, US). This apparatus is equipped with three PVC chambers, each with dimensions of 50 x 15 x 30 cm. In this procedure, mice were individually suspended by their tails, approximately 10 cm above the ground, for a period of 6 min. A critical measure in this test is the duration of immobility, widely recognized as an indicator of depression-like behavior. This parameter was meticulously monitored through strain sensors affixed to the apparatus. All data obtained during the test were digitally captured and subsequently analyzed using the BIO-TST5 software (Bioseb, US).

### 2.8 *Ex vivo* field EPSP (fEPSP) recordings

Once the behavioral tests were concluded, horizontal hippocampal slices from sham and o*Aβ*_1-42_ mice were prepared using a well-established method (Fig. 1C). In brief, animals were deeply anesthetized with halothane (Fluothane, AstraZeneca, UK) and subsequently decapitated. The brains were swiftly removed and immediately immersed in an ice-cold, oxygenated (95% O2 – 5% CO2) cutting solution. This solution comprised the following components (all in mmol/L, sourced from Sigma, US): 40 NaCl (#S9888), 10 glucose (#G8270), 150 sucrose (#84100), 1.25 NaH2PO4 (#S8282), 26 NaHCO3 (#S6014), 4 KCl (#P3911), 0.5 CaCl2 (#499609), and 7 MgCl2 (#208337). Brains were then sectioned on a vibratome (7000smz-2; Campden Instruments, UK) to produce 300 µm thick slices containing the dorsal hippocampus. These slices were incubated for over 1.5 h at room temperature (22°C) in oxygenated artificial cerebrospinal fluid (aCSF), which contained (in mmol/L): 125 NaCl (#S9888), 3.5 KCl (#P3911), 2.4 CaCl2 (#499609), 1.3 MgCl2 (#208337), 26 NaHCO3 (#S6014), 25 glucose (#G8270), and 1.2 NaH2PO4 (#S8282), all obtained from Sigma, US.

Electrophysiological recordings were performed using two setups, each consisting of a multi-electrode array (MEA2100-Mini-System) pre-amplifier and a filter amplifier, controlled by a data acquisition card and MC_Experimenter V2.20.0 software. Individual slices were placed in MEA recording chambers (MEA60; Multi Channel Systems, Reutlingen, Germany), continuously perfused with aCSF at a flow rate of 2 mL/min, and maintained at 32 °C. The MEA was situated on an inverted MEA-VMTC-1 video microscope and included 60 extracellular electrodes, spaced 200 μm apart. Each electrode functioned as either a recording or stimulation electrode. Slices were secured against the electrode array with a nylon mesh. Stimulation was delivered via one electrode in the Schaffer Collateral pathway (S1) using a stimulus generator unit, while field excitatory postsynaptic potentials (fEPSPs) were simultaneously recorded in the CA1 *stratum radiatum* from the other electrodes. A second electrode (S2) stimulated an independent pathway as a control.

After a 20-min equilibration period, baseline fEPSPs at the CA3-CA1 synapse were measured for 30 min. Veh. or PRE-084 (1 µM) was perfused 15 min into this baseline recording. Long-Term Potentiation (LTP) was induced using a high-frequency stimulation (HFS) protocol, consisting of three 100-Hz trains, each lasting 1 s, with 10-min intervals between them. Veh. or PRE-084 was discontinued 5 min after HFS and fEPSPs were monitored for 60 min post-HFS to evaluate LTP induction.

Data analysis was conducted using Multichannel Analyzer software (V2.20.0). As the responses did not include population spikes, the amplitude of fEPSPs was quantified by their peak-to-peak value. Results are presented as mean ± SEM, with ’n’ representing the number of slices analyzed. Data from three different recording electrodes were utilized for each slice.

### 2.9 Immunohistochemistry staining

To assess the distribution of S1Rs in the dorsal hippocampus, immunohistochemistry staining was conducted for illustrative purposes. After concluding all experiments, a select group of mice were anesthetized with an *ip.* injection of ketamine/xylazine (75/10 mg/kg; KETALAR, Pfizer and ROMPUM, Bayer). After anesthesia, the mice underwent transcardial perfusion with 0.9% saline followed by 4% paraformaldehyde in PBS (0.1 M, pH 7.4). The brains were then removed, cryoprotected in 30% sucrose in PB, and coronal sections (40 µm) were prepared using a sliding freezing microtome (Microm HM 450). These sections were preserved at -20°C in 50% glycerol in PBS.

For fluorescence immunohistochemistry, free-floating sections were initially incubated for 45 min in 10% normal donkey serum (NDS; RRID:AB 2810235, Sigma-Aldrich, US) in Tris-buffered saline (TBS) containing 0.1% Triton X-100 (TBS-T; #T8532, Sigma, US). This was followed by overnight incubation at room temperature with polyclonal rabbit anti-S1R primary antibody (1:200; RRID:AB_2301712, Proteintech Group, US) prepared in the same buffer with 0.05% sodium azide and 5% NDS. The following day, sections were rinsed and incubated for 2 h with FITC-conjugated donkey anti-rabbit antibody (1:150; RRID:AB_2315776, Jackson ImmunoResearch, US) in TBS-T. After several washes, the sections were treated with 0.01% DAPI (#sc-3598, Santa Cruz Biotechnology) in TBS for 5 min, then washed, mounted on gelatinized slides, dehydrated, and coverslipped using a fluorescence mounting medium (Dako, Agilent, US). Imaging was conducted using a confocal microscope (LSM 800, Carl Zeiss) at 10x and 40x magnifications.

### 2.10 Statistical analysis

In this study, data are reported as mean ± standard error of the mean (SEM). Statistical analyses were tailored to the experimental design: one-way or two-way Analysis of Variance (ANOVA) was applied, with time as a within-subjects factor and treatment as a between-subjects factor. Post hoc analyses following ANOVA were conducted using Tukey’s test for detailed comparisons. In cases involving only two groups, the Student’s t-test was utilized. The threshold for statistical significance was established at *p* < 0.05.

Statistical computations were performed using SPSS software version 24 (RRID:SCR_002865; IBM, US) and GraphPad Prism version 8.3.1 (RRID:SCR_002798; Dotmatics, US). The graphical presentation of the data, including the creation of final figures, was executed using CorelDraw X8 Software (RRID:SCR_014235; Corel Corporation, Canada).

## 3. Results

### 3.1. Activation of S1Rs restores dorsal CA1 long-term synaptic plasticity in early AD model

To provide a foundational understanding at the synaptic level before progressing to behavioral analyses, our study began by analyzing synaptic plasticity, a process notably disrupted by o*Aβ* as extensively reported [29]. Specifically, we investigated the long-term effects of a single *icv.* injection of o*Aβ*_1-42_ and the influence of S1R activation on LTP at the dorsal CA3-CA1 synapse in an *ex vivo* preparation (Fig. 2A). To achieve this, sham and o*Aβ*_1-42_ mice were sacrificed and hippocampal slices were collected.

**Fig. 2.**
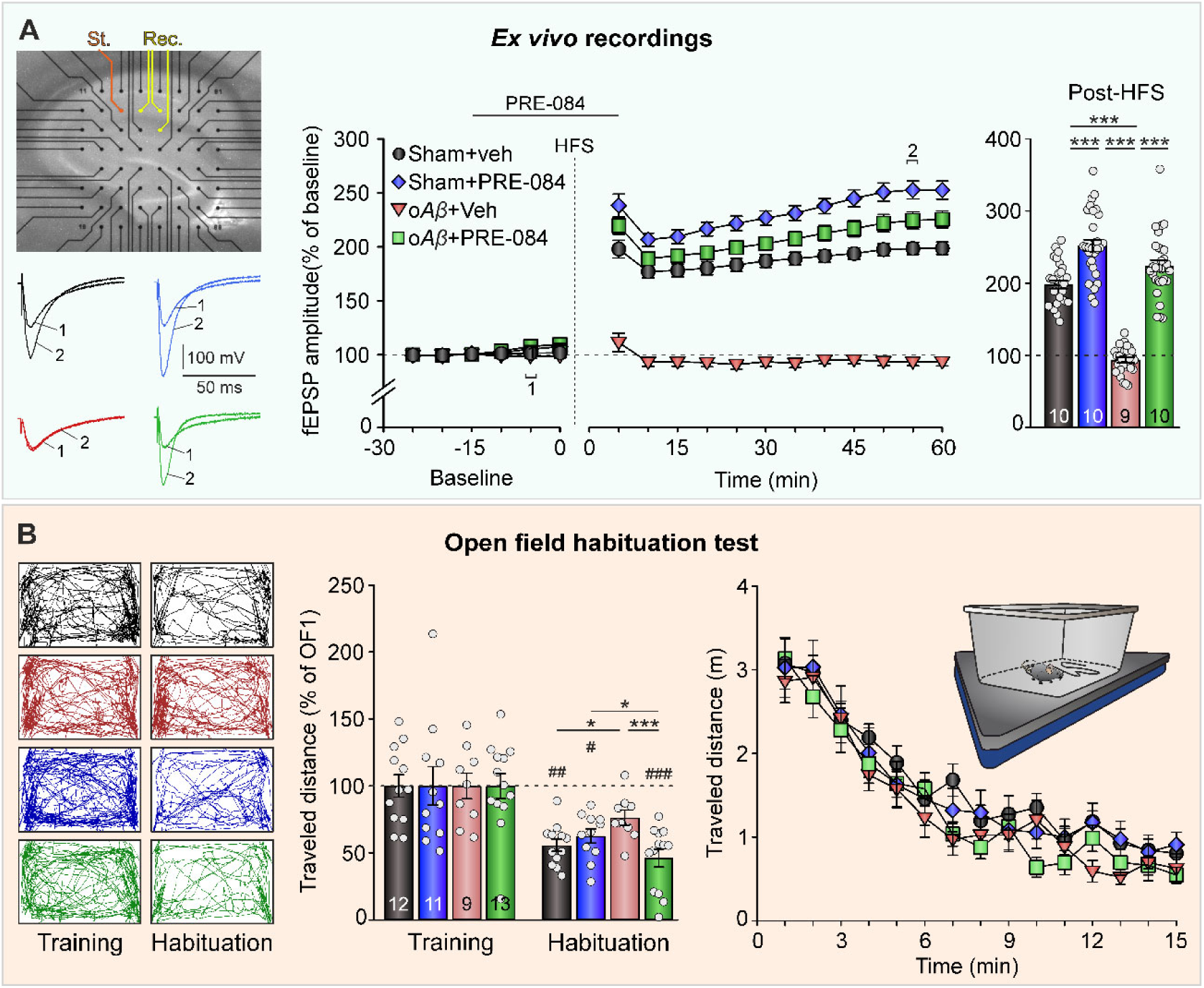
Pharmacological activation of S1Rs with PRE-084 improves LTP at the CA3-CA1 synapse and hippocampal-dependent contextual memory deficits induced by o*Aβ*_1-42_. **A.** 6 days following *icv.* infusion of veh. (PBS; sham animals) or o*Aβ*_1-42_, long-term synaptic plasticity at the CA3-CA1 synapse was examined *ex vivo*. Slices were treated with veh. (saline) or PRE-084 during the 15 min preceding and for 5 min subsequent to HFS. The amplitude of fEPSPs evoked in the CA1 region by single-pulse stimulation of the Schaffer collaterals was analyzed both pre-HFS (baseline) and post-HFS. Representative fEPSPs traces, depicted on the left, correspond to recordings before (1) and after (2) HFS, as referenced in the middle graph. This graph represents the temporal progression of fEPSP amplitude, expressed as percentage of baseline (100%). The right barplot illustrates the potentiation levels observed during final last 10 min of post-HFS recordings, with the number of slices indicated for each experimental group**. B.** 20 min after *ip.* administration of veh. or PRE-084, and 1 h after *icv.* infusion of veh. (sham) or *Aβ*_1-42_, mice underwent a 15-min training session (OF1) in the open field habituation test. A second *ip.* injection of veh. or PRE-084 o was given 24 h later, 20 min before a 15-min habituation session (OF2; recall trial). The left panels display LABORAS®-generated trackings of mice movements during OF1 and OF2 sessions. Middle bar plots quantify the distances travelled during both OF1 and OF2, expressed as a percentage of OF1 (100%). The right graph shows the minute-by-minute travel distances during the OF1 session. The number of animals of each experimental group is indicated in the corresponding bars. Data are represented as mean ± SEM. \**p* < 0.05, ** *p* < 0.01, *** *p* < 0.001, intergroup differences. # *p* < 0.05, ## *p* < 0.01, ### *p* < 0.001, intragroup differences (*vs*. training).

Electrical single-pulse stimulation was applied at the Schaffer collateral pathway to evoke a fEPSP at the CA1 *stratum radiatum*. The amplitude of this synaptic response was monitored both before and after LTP induction to assess potential changes in synaptic transmission. Prior to LTP induction, the perfusion of veh. (veh._(pf.)_) did not alter the basal fEPSP amplitude in slices from sham (n = 4 animals, 10 slices; F_(2.673, 72.183)_ = 1.529, *p* = 0.218) or o*Aβ*_1-42_ (n = 4 animals, 9 slices; F_(2.542, 66.091)_ = 0.629, *p* = 0.573) mice. In contrast, the perfusion of PRE-084 significantly increased the basal fEPSP amplitude in slices from both sham (n = 4 animals, 10 slices; F_(2.095, 60.744)_ = 14.372, *p* < 0.001) and o*Aβ*_1-42_ (n = 4 animals, 10 slices; F_(3.129, 90.740)_ = 48.003, *p* < 0.001) mice. Upon baseline establishment, the application of a HFS protocol induced robust synaptic potentiation in slices treated with sham+veh._(pf.)_ (189.8 ± 5.5% of baseline; F_(1.826, 49.298)_ = 188.506, *p* < 0.001) and sham+PRE-084_(pf.)_ (234.4 ± 7.1% of baseline; F_(1.686, 48.896)_ = 257.792, *p* < 0.001), which remained stable for at least 1 h in these conditions. Notably, the magnitude of potentiation was significantly higher in slices treated with sham+PRE-084_(pf.)_ compared to those receiving sham+veh. _(pf.)_ (*p* < 0.001). In contrast, slices treated with o*Aβ*_1-42_+veh._(pf.)_ showed no potentiation (93.5 ± 3.6% of baseline; F_(2.593, 67.415)_ = 1.926, *p* = 0.142). Yet, when PRE-084 was perfused in slices from o*Aβ*_1-42_-treated mice, LTP was successfully restored (o*Aβ*_1-42_+PRE-084_(pf.)_; 209.8 ± 6.7% of baseline; F_(1.609, 46.650)_ = 193.344, *p* < 0.001). Significant differences were observed between o*Aβ*_1-42_+veh._(pf.)_ group and the other three conditions, even 50 min after HFS application (F_(3,111)_ = 99.506, *p* < 0.001; *post hoc vs*. sham+veh._(pf.)_, *p* < 0.001; *post hoc vs*. sham+PRE-084_(pf.)_, *p* < 0.001; *post hoc vs*. o*Aβ*_1-42_+PRE-084_(pf.)_, *p* < 0.001).

### 3.2 S1R activation improves habituation memory deficits induced by acute hippocampal amyloidosis

The electrophysiological results indicate a role for S1R in regulating synaptic plasticity at the dorsal CA3-CA1 synapse, a crucial region for processing both non-associative and associative memory forms, including contextual memory [30–32]. Consequently, the effects of the pharmacological S1R activation on habituation memory, a non-associative hippocampal-dependent learning form [25], were examined using the OF habituation test in the automated LABORAS (Fig. 2B). In the first session of the task, all experimental groups demonstrated a similar and progressive reduction in the distance traveled (sham+veh._(ip.)_, n = 12, F_(14,154)_ = 28.218, *p* < 0.001; sham+PRE-084_(ip.)_, n = 11, F_(14,140)_ = 24.260, *p* < 0.001; o*Aβ*_1-42(_+veh._(ip.)_, n = 9, F_(14, 112)_ = 24.805, *p* < 0.001; o*Aβ*_1-42_+PRE-084_(ip.)_, n = 13, F_(14, 168)_ = 39.779, *p* < 0.001), suggesting intact contextual memory encoding. However, upon reexposure to the same platform the following day, significant reductions in locomotor activity were observed in mice treated with sham+veh._(ip.)_ (t(11) = 3.911, *p* = 0.002) sham +PRE-084_(ip.)_ (t(10) = 2.259, *p* = 0.047) and o*Aβ*_1-42_+PRE-084_(ip.)_ (t(12) = 6.946, *p* < 0.001), but not in those receiving o*Aβ*_1-42_+veh._(ip.)_ (t(8) = 1.874, *p* = 0.098). The distance traveled by the o*Aβ*_1-42_+veh._(ip.)_ group was significantly higher (F_(3, 44)_ = 5.42, *p* = 0.003) than that of the sham+veh._(ip.)_ (*post hoc vs*. o*Aβ*_1-42_+veh._(ip.)_, *p* = 0.017) or o*Aβ*_1-42_+PRE-084 (*post hoc vs*. o*Aβ*_1-42_+veh._(ip.)_, *p* < 0.001) groups, indicating impaired habituation memory retrieval. Noteworthy is the observation that animals injected with o*Aβ*_1-42_+PRE-084_(ip.)_ exhibited more robust habituation between sessions compared to those treated with sham+PRE-084_(ip.)_ (*p* = 0.009).

### 3.3 Social memory capabilities are improved by S1R activation in the early AD model

In addition to its established role in spatial memory forms, the dorsal pole of hippocampus, particularly the CA2 region, has also been identified as essential for social memory [33, 34], a cognitive function significantly impacted in AD [35], yet not extensively studied. To address this gap and considering the widespread expression of S1R in the dorsal hippocampus including the CA2 region, as shown by immunohistochemical staining (Fig. 3A), we aimed to elucidate the impact of o*Aβ*_1-42_ and the potential modulatory effects of S1R pharmacological activation on social memory in our amyloidosis model. Accordingly, the social recognition test was employed (Fig. 3B,C). During the initial training phase, no significant difference was observed in the time spent exploring the empty cylinder compared to an unknown conspecific across all treatment groups (sham+veh._(ip.)_, n = 9, t(8) = 0.988, *p* = 0.352; sham+ PRE-084, n = 8, t(7) = 0.297, *p* = 0.775; o*Aβ*_1-42_+veh._(ip.)_, n = 7, t(6) = 1.407, *p* = 0.209; o*Aβ*_1-42_+PRE-084_(ip.)_, n = 7, t(6) = 0.659, *p* = 0.534). Furthermore, no treatment-related differences were observed in the time exploring either the empty cylinder (F_(3,30)_ = 0.288, *p* = 0.834) or the subject (F_(3,30)_ = 1.538, *p* = 0.227), indicating unaffected sociability. However, in the recall phase, a disparity emerged. Mice injected with sham+veh._(ip.)_ (t(8) = 3.718, *p* = 0.006) or o*Aβ*_1-42_+PRE-084_(ip.)_ (t(6) = 2.936, *p* = 0.026) demonstrated increased time exploring an unknown subject compared to a familiar conspecific. In contrast, the groups receiving sham+PRE-084_(ip.)_ (t(7) = 1.465, p = 0.186) or o*Aβ*_1-42_+veh._(ip.)_ (t(6) = 1.960, *p* = 0.098) showed similar exploration times for both subjects. This pattern suggests that S1R pharmacological activation induces social memory deficits in healthy subjects but rescues social memory in those undergoing acute hippocampal amyloidosis.

**Fig. 3.**
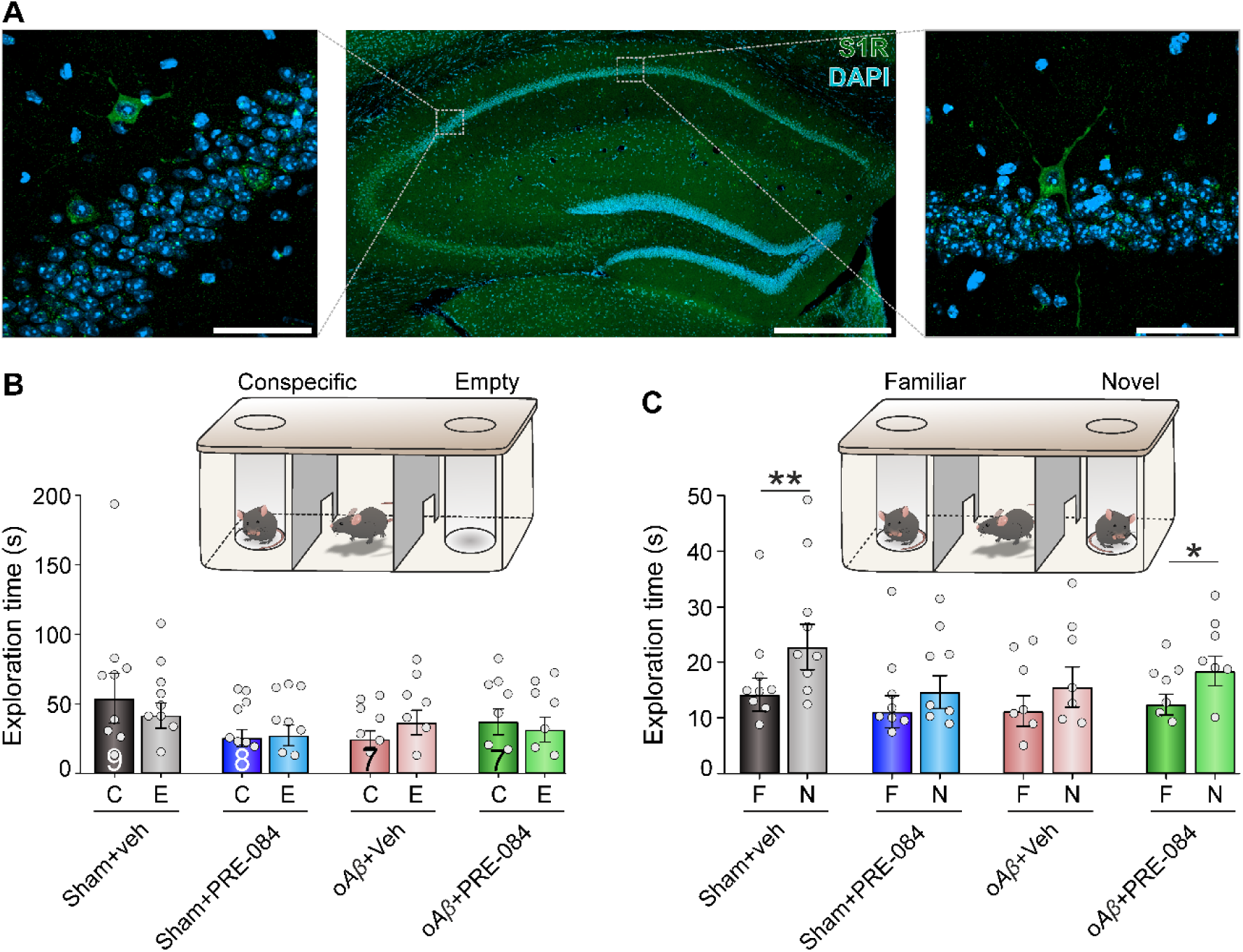
Pharmacological activation of S1Rs in the dorsal hippocampus with PRE-084 counteracts o*Aβ*_1-42_-induced social memory deficits. **A**. Confocal immunofluorescence images of 40-μm thick coronal sections stained with anti-S1R antibody (green) and counterstained with DAPI (blue), illustrating dense S1R expression throughout the hippocampus, including detailed views of the CA2 (left inset) and CA1 (right inset) regions. Scale bar, 500 μm (main image) and 50 μm (insets). **B.** Social memory test.1 h after *icv.* drug administration and 20 min following *ip.* injections, mice were introduced to an arena containing two empty cylinders for a 5-min habituation period. Subsequently, an unknown conspecific was placed in one of the cylinders, and the experimental subject was allowed 10 min of free exploration during the training phase. The exploration times for the conspecific (C) and the empty cylinder (E) were recorded and quantified. **C.** 24 h after the training phase, a 5-min recall session was conducted, where the familiar animal (F) from the training and a novel mouse (N) were placed in separate cylinders. The duration of interaction of the experimental subject with each stimulus mouse was measured. The number of animals in each experimental group is indicated on the corresponding bar. Data are represented as mean ± SEM. \**p*<0.05, \*\**p*<0.01.

### 3.4 S1R activation in dorsal hippocampus does not alter motor function and does not induce in anxiety-nor depression-like behaviors

To rule out the possibility that the interpretation of data from learning and memory tests is influenced by motor impairments or emotional disturbances caused by the administered compounds, a series of behavioral tests were conducted (Fig. 4). These tests aimed to evaluate motor function as well as anxiety- and depression-related behaviors. Initially, locomotor activity in the training session of the open field habituation test was assessed using the automated LABORAS (Fig. 4B). The distance traveled by all experimental groups (sham+veh._(ip.)_, n = 12; sham+PRE-084_(ip.)_, n = 11; o*Aβ*_1-42_+veh._(ip.)_, n = 9; o*Aβ*_1-42_+PRE-084_(ip.)_, n = 13) was similar (F_(3,41)_ = 0.694, *p* = 0.561), indicating that motor function was not compromised. Additionally, LABORAS quantified the time mice spent on various predefined spontaneous behaviors (sham+veh._(ip.)_, n = 11; sham+PRE-084_(ip.)_, n = 10; o*Aβ*_1-42_+veh._(ip.)_, n = 9; o*Aβ*_1-42_+PRE-084_(ip.)_, n = 12). No significant differences were detected across the groups for behaviors such as locomotion (F_(3,41)_ = 0.442, *p* = 0.725), climbing (F_(3,41)_ = 0.112, *p* = 0.953), rearing (F_(3,41)_ = 0.755, *p* = 0.527) and grooming (F_(3,41)_ = 0.815, *p* = 0.493; Fig. 4B). These findings suggest that motor functions, as well as stress or anxiety levels, were unaffected by the treatment. The elevated plus maze was employed to further assess locomotor activity and anxiety-like behaviors (sham+veh._(ip.)_, n = 13; sham+PRE-084_(ip.)_, n = 9; o*Aβ*_1-42_+veh._(ip.)_, n = 9; o*Aβ*_1-42_+PRE-084_(ip.)_, n = 11; Fig. 4A).

**Fig. 4.**
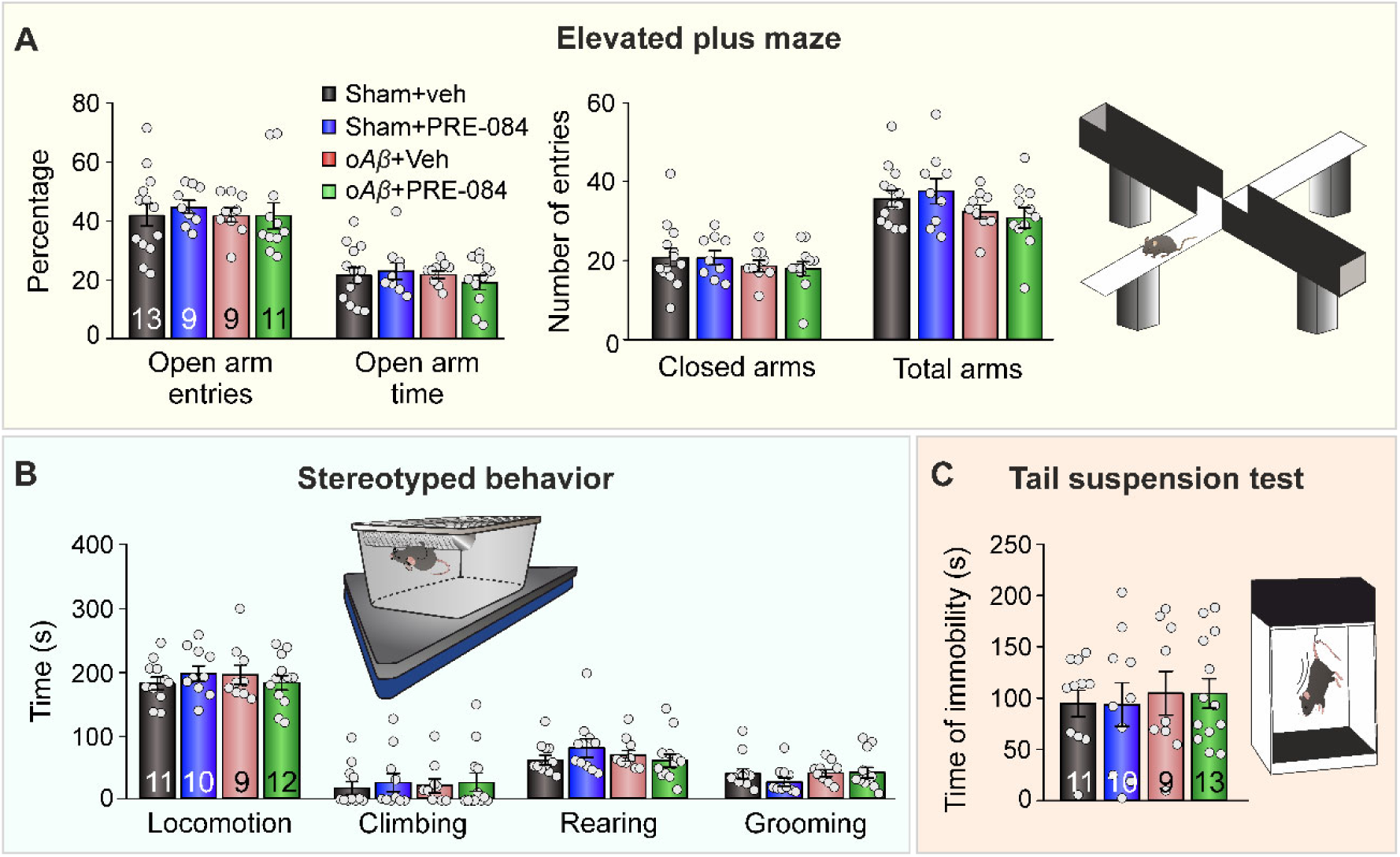
Intraperitoneal administration of PRE-084, alone or in combination with *icv.* injections of o*Aβ*_1-42_, does not induce alterations of motor function nor anxiety or depression-like behaviors. Different behavioral tests were performed 20 min (stereotyped behavior, tail suspension test) or 35 min (elevated plus maze) after *ip.* injections of veh. (saline) or PRE-084. These tests occurred 48 h (stereotyped behavior, elevated plus maze) or 72 h (tail suspension test) following *icv.* infusions of veh. (PBS; sham animals) or o*Aβ*_1-42_. **A.** Elevated plus maze. Mice were exposed to the platform for a 5-min session. The left graph displays the percentage of open arms entries, and the time spent in open arms. The right bar plots show the number of entries into closed arms and the total number of entries across all arms. **B.** Stereotyped behavior. This assessment measured the time spent in various behaviors was measured during a 15-min session using the LABORAS^®^, which detects vibration patterns generated by the animals’ movements. **C.** Tail suspension test. The duration of immobility was quantified over a 6-min session, indicating potential depression-like behavior. The number of animals in each experimental group is indicated on the corresponding bar. Data are represented as mean ± SEM.

No significant differences were found between groups in terms of the number of entries into closed arms (F_(3,41)_ = 0.523, *p* = 0.669) and total arms (F_(3,41)_ = 1.524, *p* = 0.224), supporting the notion that motor function remained intact. Moreover, all experimental groups exhibited similar percentages of open arm entries (F_(3,41)_ = 1.165, *p* = 0.336) and time spent in open arms (F_(3,41)_ = 0.416, *p* = 0.743), indicating no anxiety-related behaviors due to the treatment. Lastly, the tail suspension test was used to measure immobility, serving as an indicator of potential depression-related behaviors (Fig. 4C).

No differences were observed between groups in this regard (sham+veh._(ip.)_, n = 11; sham+PRE-084_(ip.)_, n = 10; o*Aβ*_1-42_+veh._(ip.)_, n = 9; o*Aβ*_1-42_+PRE-084_(ip.)_, n = 13) (F_(3,42)_ = 0.112, p = 0.946). Collectively, these results provide substantial evidence that pharmacological S1R activation in the dorsal hippocampus, both in healthy individuals and in a scenario of acute hippocampal amyloidosis, does not induce mood disturbances.

## 4. Discussion

In the early stages of AD, a notable imbalance in neuronal activity across various brain regions is observed, with hippocampal dysfunction being particularly pronounced [3]. This dysfunction, extensively documented in both human patients and animal models [36], is largely attributed to the deleterious impact of soluble oligomeric forms of *Aβ* [37]. These oligomers induce hippocampal hyperexcitability and contributes to cognitive impairments, marking the initial stages of AD [2, 38, 39]. The increasing awareness of these early neuronal changes has spurred a concerted effort to identify therapeutic targets capable of mitigating these alterations before they culminate in irreversible neuronal damage [1].

Our research group has meticulously characterized the effects of acute amyloidosis generated by oA*β*_1-42_ on hippocampal functionality, documenting increases in neuronal excitability, alterations in synaptic plasticity, aberrant oscillatory activity, and sustained deficits in hippocampus-dependent memory [6–8, 40]. While GIRK channel activation has been proposed to control hippocampal excitability in early amyloidosis, yielding promising results [7–9], the intricacies of this channel’s activity under physiological conditions [27] and the complex nature of amyloid pathology underscores the need for additional therapeutic avenues. In this line, recent genetic and pharmacological research has highlighted the S1R as a potential target in neurodegenerative diseases, especially AD [10]. Additionally, the newly described direct relationship between S1R function and GIRK channel activity [41] positions S1R as a promising candidate to extend our previous research and explore its therapeutic potential in this pathology. In this work, we evaluated the efficacy of S1R activation in reversing synaptic and cognitive dysfunctions induced by o*Aβ*_1-42_ using an early hippocampal amyloidosis model in mice of both sexes, and assessed its influence on social memory, a less frequently explored aspect in this pathology. Given that, as previously described, our model shows the same effects regardless of sex [6] and as no differences between sexes were detected in this particular study (data not shown), data from male and female mice were combined to increase the sample size and obtain more robust results.

Electrophysiologically, our observations revealed that, beginning 7 days post a single *icv.* injection of o*Aβ*_1-42_, hippocampal slices from these mice failed to exhibit LTP at the CA3-CA1 synapse following a HFS protocol. This aligns with our previous findings [6] and corroborates the well-documented fact that soluble o*Aβ* disrupts synaptic plasticity in the hippocampus [42]. Remarkably, perfusion of the slices with PRE-084, a selective S1R agonist [43], resulted in an increase of the amplitude of baseline fEPSPs pre-HFS in both control and o*Aβ*_1-42_-treated animals, reflecting the S1R’s capacity to modulate glutamatergic transmission in the hippocampus [17]. Notably, following HFS application, S1R activation by PRE-084 was able to rectify the LTP impairment caused by o*Aβ*. This observation not only aligns with findings from other authors [44] but also expands upon them by illustrating the potential of S1R activation to reverse the lasting LTP alterations resulting from initial o*Aβ*_1-42_ exposure, as evidenced in slices obtained several days after a single *icv.* injection of the peptide. The interaction of S1Rs with various ion channels grants them considerable influence over neuronal excitability and synaptic plasticity [45], suggesting their activation could counteract *Aβ*-induced hyperexcitability, thereby reinstating the generation of LTP [39]. One such interaction is with GIRK channels, directly associated with S1R function [41], which activation has been shown to counteract the negative effects of o*Aβ*_1-42_ on LTP *ex vivo* and *in vivo* [7, 8, 40]. Additional mechanisms, such as the enhancement of NMDA receptor-mediated trafficking and activity [46], or upregulation of synaptic plasticity regulators like BDNF [47], further reinforce the beneficial impact of S1Rs on synaptic plasticity processes. Interestingly, the administration of PRE-084 in control slices also augmented LTP, very likely by same mechanism that increased basal EPSPs amplitude through preventing a small-conductance Ca^2+^-activated K^+^ current (SK channels) and enhancing NMDAR responses and LTP [17], underscoring the potential of S1Rs not only in ameliorating deficits but also in boosting hippocampal synaptic plasticity under normal conditions.

Next, we aimed to elucidate how these findings on synaptic plasticity of the CA3-CA1 pathway might influence memory and learning processes. The anti-amnesic effects of S1Rs, which affect various phases of memory processing across different types of hippocampus-dependent memory, have been long recognized [48–50]. Here, to assess hippocampal-dependent contextual memory, we employed an open field habituation test [25], focusing particularly on two crucial aspects of memory processing: encoding and retrieval [51, 52]. All experimental groups exhibited a gradual increase in memory formation during the initial session, indicative of successful memory encoding. However, 24 h post-injections, the o*Aβ*_1-42_-treated group displayed significant deficits in retrieving habituation memory. These observations are in line with prior research documenting hippocampal-dependent memory impairments in early amyloidosis models [6–8]. Significantly, PRE-084 administration mitigated these deficits, echoing research that shows that S1R agonists alleviate learning impairments induced by *icv.* injections of *Aβ*_25-35_, the biologically active isoform of the peptide [53], in various memory tasks [24]. The broad effectiveness of S1R activation in alleviating hippocampus-dependent cognitive alterations, as evidenced in spatial memory tasks in transgenic AD models [54, 55] and models involving *icv.* injections of *Aβ*_1-42_ [56], highlights its therapeutic potential. Moreover, these results underscore that the positive effects of activating S1Rs on hippocampal synaptic plasticity extend to ameliorating cognitive deficits associated with it.

Subsequently, we extended our research scope to evaluate social memory, an aspect that has been less examined in neurodegenerative disease studies. Social memory is closely associated with hippocampal functionality, particularly involving neurons in the CA2 region [57], with recent evidence pointing to the significance of CA2-CA1 neuronal communication in long-term memory formation [58, 59]. Considering the observed alterations in hippocampal neuronal communication in our amyloidosis model, and given the distribution of S1Rs across the CA3-CA1 region including CA2, as evidenced here and elsewhere [60], we hypothesized that social memory might be compromised. Indeed, our findings confirmed that mice administered with o*Aβ*_1-42_, unlike those treated with veh. (sham), exhibited impaired ability to differentiate between novel and familiar counterparts, indicative of disrupted social memory. This finding is consistent with prior studies showing that social recognition memory is affected in transgenic disease models [61, 62] and by intrahippocampal injections of *Aβ*_1-42_ [63]. Once again, administration of PRE-084 mitigated these memory deficits, reinforcing its potential role in rectifying cognitive deficits associated with hippocampal function due to amyloidosis. However, it is noteworthy that, unlike its effect on habituation memory, PRE-084 administration in control mice impacted this type of memory. This discrepancy underscores the necessity for more in-depth studies into the role of S1Rs in the mechanisms governing social memory and highlights the importance of dose optimization for the agonist to mitigate potential adverse effects.

Importantly, our observations confirmed the absence of behaviors indicative of motor, anxiety, or depressive disorders in the experimental animals. This suggests that the observed memory alterations were indeed cognitive in nature, rather than secondary to other behavioral impairments. Additionally, these results highlight the potential safety of PRE-084 for therapeutic application, emphasizing the importance for treatments that do not adversely affect motor functions or mood, especially given the complex nature of the brain’s responses to pharmacological interventions.

The stable expression of S1Rs during normal human aging, along with their potential upregulation [64, 65], identifies them as viable targets for addressing age-related cognitive decline. Furthermore, the reduced expression of S1Rs in AD patients [66] underscores their potential role in the defense mechanisms against neurodegenerative conditions. The neuroprotective properties of PRE-084, evident in cultures treated with *Aβ*_25-35_ [67] and various *in vivo* pharmacological models [68, 69], provide further support to this hypothesis. The exploration of S1R agonists in the treatment of other neurodegenerative disorders, including Parkinson’s disease and multiple sclerosis, points to their wider therapeutic relevance [70]. Interestingly, Donepezil, a medication commonly prescribed in AD management, shows a high affinity for S1Rs [70]. However, its effects are primarily symptomatic, without halting the progression of the disease [71]. This insight advocates for the development of multi-target therapeutic strategies that combine S1R agonists with other synergistic drugs. Current research in this area is still in its early stages [72]. Such strategies should aim to counteract the initial hyperexcitability triggered by *Aβ* [39], leveraging targets such as S1Rs and GIRK channels, as previously analyzed [7, 8], and promote the removal of the peptide to prevent further neuronal damage. The latter approach has been the focus of extensive recent investigations, yielding promising outcomes [1, 73]. Such integrated approached could pave the way for more comprehensive treatments for neurodegenerative diseases.

## 5. Conclusions

In conclusion, our study provides compelling evidence for the potential of S1R activation in reversing synaptic and cognitive deficits induced by o*Aβ* in early amyloidosis, applicable to mice of both sexes. The administration of the S1R agonist, PRE-084, not only mitigated the o*Aβ*-induced persistent abnormalities in LTP but also effectively restored hippocampus-dependent memory functionalities. Notably, our findings underscore the therapeutic potential of S1R activation in addressing complex cognitive processes, with a particular emphasis on social memory—a facet critically impacted yet underexplored in neurodegenerative diseases [74]. The absence of adverse behavioral outcomes associated with PRE-084 treatment accentuates its safety profile, underscoring its potential as a therapeutic agent. Moving forward, we advocate for comprehensive research that incorporates multitarget strategies including S1R. Prioritizing dose optimization and unraveling the detailed molecular mechanisms behind S1R’s neuroprotective effects are essential steps before broadening its therapeutic applications. This approach not only promises to enhance our understanding of AD treatment but also opens doors to innovative, more effective interventions for this challenging neurodegenerative condition.

## Declarations

### Data availability statement

All data supporting the results in the paper are in the paper itself and the supporting information. Raw data and recordings are available in the laboratory from the corresponding authors and will be provided upon reasonable request.

### Ethics approval

All experimental procedures were reviewed and approved by the Ethical Committee for Use of Laboratory Animals of the University of Castilla-La Mancha (PR-2021-12-21 and PR-2023-25) and conducted according to the European Union guidelines (2010/63/EU) and the Spanish regulations for the use of laboratory animals in chronic experiments (RD 53/2013 on the care of experimental animals: BOE 08/02/2013).

### CRediT authorship contribution statement

LJD and JDNL were responsible for the initial conceptualization; GIL, RJH, and SD performed methodology, data curation and formal analysis; SD and RJH were responsible for the visualization; SD for writing the original draft; JDNL, SD and LJD did the writing – review and editing. LJD and JDNL were responsible for funding acquisition, project administration and supervision. All authors read and approved the final manuscript.

